# How to use online tools to generate new hypotheses for mammary gland biology research: a case study for *Wnt7b*

**DOI:** 10.1101/2020.09.19.304667

**Authors:** Yorick Bernardus Cornelis van de Grift, Nika Heijmans, Renée van Amerongen

**Author notes:** Twitter: @wntlab (Renée van Amerongen).

## Abstract

An increasing number of ‘-omics’ datasets, generated by labs all across the world, are becoming available. They contain a wealth of data that are largely unexplored. Not every scientist, however, will have access to the required resources and expertise to analyze such data from scratch. Luckily, a growing number of investigators is dedicating their time and effort to the development of user friendly, online applications that allow researchers to use and investigate these datasets. Here, we will illustrate the usefulness of such an approach.

Using regulation of *Wnt7b* as an example, we will highlight a selection of accessible tools and resources that are available to researchers in the area of mammary gland biology. We show how they can be used for *in silico* analyses of gene regulatory mechanisms, resulting in new hypotheses and providing leads for experimental follow up. We also call out to the mammary gland community to join forces in a coordinated effort to generate and share additional tissue-specific ‘-omics’ datasets and thereby expand the *in silico* toolbox.

## Introduction

The experimental technology that allows genome wide analyses at the molecular level (genomics, epigenomics, transcriptomics, metabolomics and proteomics – hereafter combinedly referred to as ‘omics’ approaches) continues to evolve at breathtaking speed. Despite the fact that these techniques are becoming more affordable and therefore more widely available for scientists worldwide, they are still quite expensive – a prohibitory factor for those with limited financial resources. This is especially true for sophisticated approaches such as single-cell RNA sequencing (scRNAseq) and other-single cell approaches that are still being developed. Moreover, not everyone will have local access to the required infrastructure. Of course, scientific collaborations can offer a solution. Even then, it can be a challenge to integrate a variety of these technologies into one’s research program [1].

As can be gleaned from the published literature, all too frequently only a few hits or top candidates are followed up in instances where genome-wide datasets are generated. As a consequence, a wealth of data remains unexplored. These datasets constitute a rich and valuable resource for the larger scientific community. As an example, we have previously used published microarray data to identify the most stably rather than the most differentially expressed genes, resulting in a new set of reference genes for qRT-PCR studies in the developing mouse mammary gland [2].

Most ‘omics’ datasets are deposited in public repositories such as the NCBI Gene Expression Omnibus (https://www.ncbi.nlm.nih.gov/geo/), either in raw format or in a more processed form. While this makes them available to all scientists in theory, in practice not everyone has the bioinformatics skills and expertise to analyze these data from scratch. Fortunately, multiple labs are dedicating their time and effort to the development of online tools that allow easy and intuitive access to these datasets, allowing researchers to explore them from the comfort of their own (home) office via a user friendly graphical interface.

Here we will highlight a selection of these online tools and demonstrate how they can be used to generate hypotheses and answer biological questions in the context of mammary gland biology. To illustrate this approach, we will build a case study around *Wnt7b*, a gene that has been implicated in mammary gland development and breast cancer, but whose precise activity and mode of regulation remain unknown.

We assume that the reader is familiar with the basic principles behind the different techniques (e.g. scRNAseq, snATACseq, Hi-C), as well as with the way in which these data are commonly presented (e.g. tSNE plots). Please note that for all figures we have kept the exact style and color schemes as generated by the different online tools to aid the reader in recognizing the output when they try out these tools for themselves.

## WNT7B in mammary gland development and breast cancer

*WNT7B* is expressed in human breast tissue and its expression has been reported to be altered in breast cancer [3,4]. Its overexpression has been associated with a poor prognosis and reduced overall survival of breast cancer patients [5]. In breast cancer, *WNT7B* has not only been shown to be expressed by the tumor cells, but also by myeloid cells present in the local microenvironment. The latter promotes angiogenesis, invasion and metastasis [6].

Its murine counterpart, *Wnt7b*, is expressed in the ductal epithelium of the mouse mammary gland [7]. The levels of *Wnt7b* remain unaltered following ovariectomy, suggesting that *Wnt7b* gene regulation is estrogen and progesterone independent [7]. During puberty, expression of *Wnt7b* is enriched in the terminal end bud epithelium, suggesting a role in branching morphogenesis [8]. *Wnt7b* has been reported to have mild transforming activities *in vitro* [9,10] and *in vivo* [11] although not all studies agree on the extent of this effect [10,12].

The precise role and regulation of *Wnt7b*/*WNT7B* in the mammary gland or breast remain unknown. So far, evidence that WNT7B protein can promote the activation of CTNNB1/TCF transcriptional complexes is lacking, despite the fact that *Wnt7b* is readily detected and shows prominent expression in luminal cells [13]. This is in contrast to other tissues, such as the skin, where the activities of WNT7B have been linked to CTNNB1/TCF driven processes [14].

## Exploring spatiotemporal patterns of *Wnt7b* expression using scRNAseq data

Public scRNAseq datasets are an ideal platform to start investigating spatiotemporal gene expression in the mammary gland [15,16]. We want to highlight two user friendly scRNAseq tools that allow analysis of the *in vivo* expression patterns of a gene of interest in both the embryonic and postnatal stages of mouse mammary gland development (Box 1). Their combined use reveals extensive details about the expression pattern of any given gene across different stages and cell populations.

**Box 1: online scRNA-seq visualization tools**

*https://marionilab.cruk.cam.ac.uk/mammaryGland/ (Bach et al*., *2017, Nature Communications* [15])

scRNAseq dataset that contains EPCAM+ sorted cells from multiple stages of the adult mammary gland cycle: (nulliparous (8w), gestation (14.5d), lactation (6d) and post-involution (11d). Expression of a gene of interest can be investigated in the context of an inferred cell type or developmental stage. Results are visualized as a tSNE plot and a box plot, both illustrating gene expression by cluster. Gene expression can also be displayed along the (luminal) differentiation trajectory in pseudotime.

*https://tabula-muris.ds.czbiohub.org/ (The Tabula Muris Consortium, 2018, Nature* [16])

Large compendium of single cell transcriptome data from the model organism *Mus musculus* that contains scRNAseq datasets from 20 adult organs and tissues, including the mammary gland. This is the only online dataset available for the mammary gland that explicitly includes stromal cells and other cell types from the supportive tissue (e.g. endothelial and immune cells). Of note, all tissues have been processed and analysed by two different protocols: cells were either FACS sorted, or single-cell sorted using microfluidic droplet-capture techniques (used for fig 1) and thus sequenced using two different methodologies, providing an innate technical validation of the data when using this tool. Fat pads 2,3 and 4 were processed from virgin mice ((10-15w), and subpopulations were separated by FACS by the following markers: Basal population (CD45^-^, CD31^-^, TER119^-^, CD49f^high-med^, CD29^med-low^), Luminal cells (CD45^-^, CD31^-^, TER119^-^, CD49f^med-low^, CD24^high-med^), mammary repopulating cells (CD45^-^, CD31^-^, TER119^-^, CD49f^high^, CD24^med^), and stromal cells (CD45^-^, CD31^-^, TER119^-^,CD49f^-^, CD24^-^).

**Figure 1.**
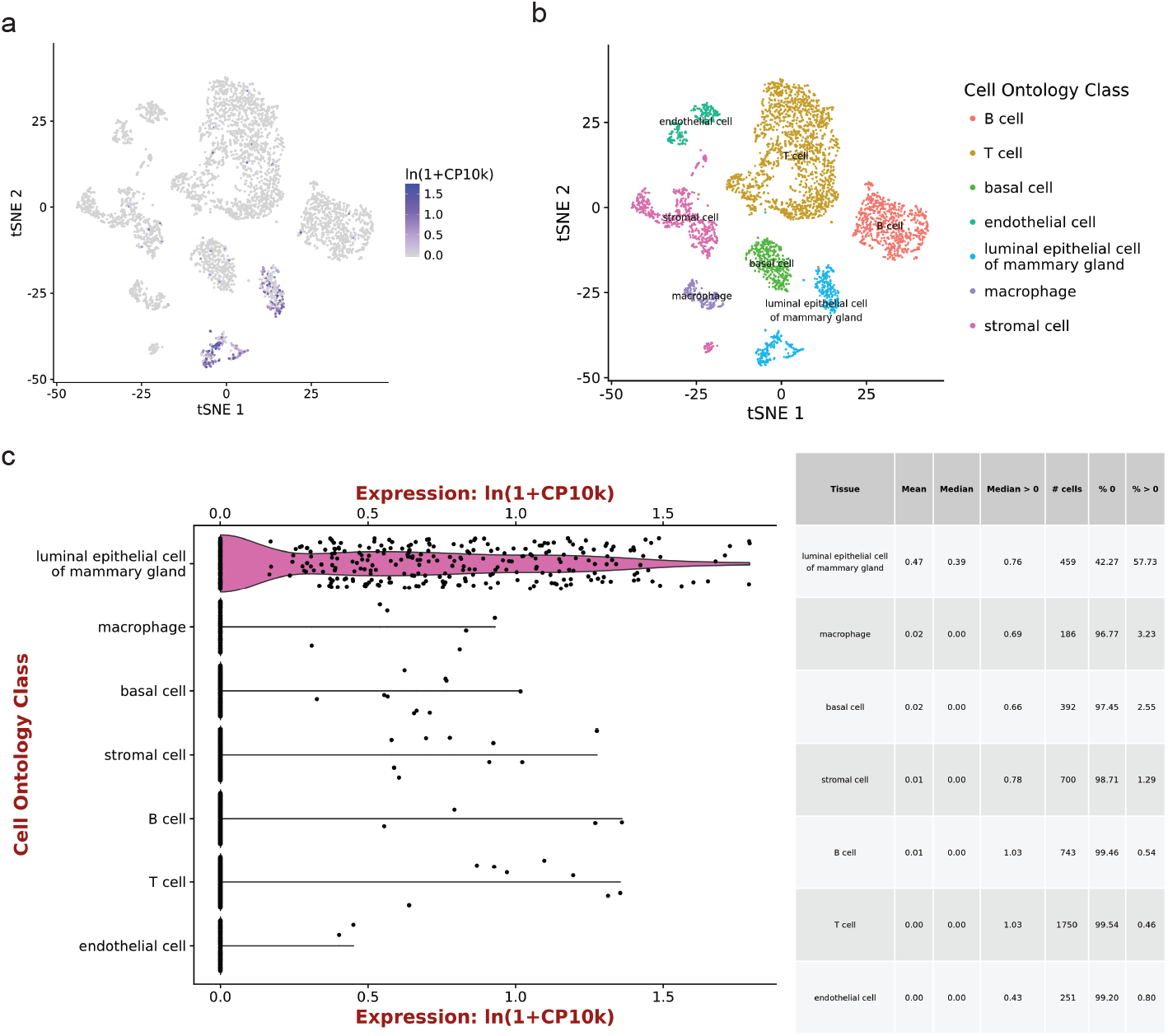
Single cell RNAseq (scRNAseq) of *Wnt7b* gene expression for all cell types in the mammary gland by Tabula Muris. A) tSNE plot displaying single cell *Wnt7b* gene expression in virgin mice superimposed on pre-defined cell clusters. Gene expression is normalised to 10,000 counts per cell. B) tSNE plot defining cell ontology of the cell clusters in A). C) Violin plot of *Wnt7b* gene expression in individual cells in the clusters defined in B). Gene expression is normalised to 10.000 counts per cell. Further relevant statistical values for each subpopulation are displayed in a table format. All plots were generated at https://tabula-muris.ds.czbiohub.org

*Wnt7b* expression is absent (or at least below the limit of detection) in the fetal mammary gland (E18, fig1a), but emerges postnatally (fig1b,c, fig2a). Its expression is cell type specific, displaying high gene expression in the luminal compartment, and low or absent expression in basal cells and supportive tissues (fat, endothelial, immune and stromal cells) (fig1c).

**Figure 2.**
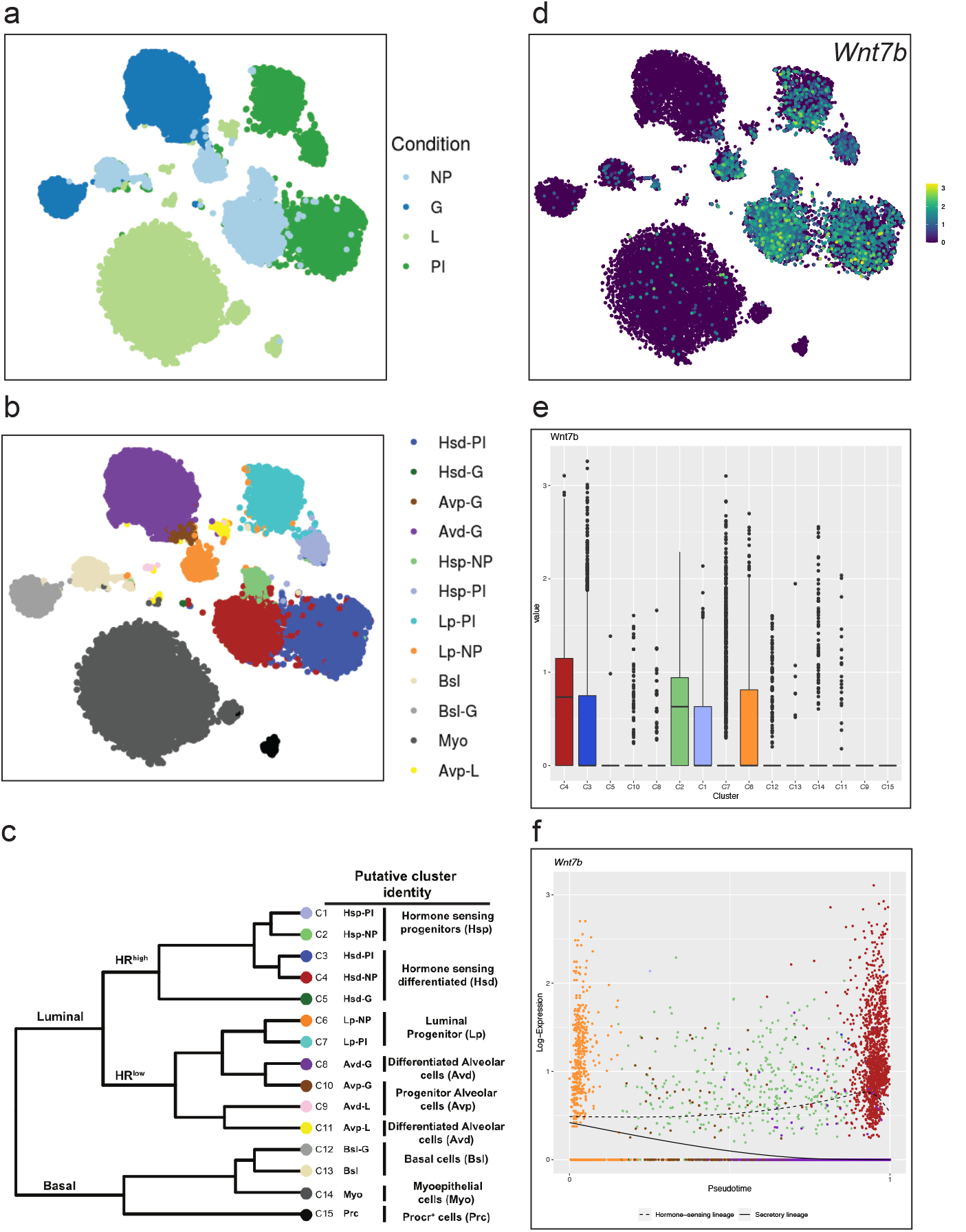
Single cell RNAseq (scRNAseq) of *Wnt7b* gene expression throughout mammary gland development A) tSNE plot displaying mammary gland developmental timepoints superimposed on pre-defined cell clusters. NP: Nulliparous, G: Gestation, L: Lactation, PI: Post-involution. B) tSNE plot defining cell ontology (through known marker genes) of cell clusters depicted in in A) & D). See C) for cell type classification. C) Dendrogram of clusters based on log transformed mean expression of 15 clusters. The tree was generated by Spearman’s rank correlation with Ward linkage. D) tSNE plot of single cell *Wnt7b* gene log transformed mean expression superimposed on pre-defined clusters. E) Bargraphs of log transformed mean expression for each 15 clusters. F) Pseudotime trajectory of the single cell Wnt7b log transformed mean expression in the luminal lineage, displaying both the average expression in the hormone sensing and secretory lineages. Each dot represents an individual cell, and the color its associated cluster. Plots were generated at https://marionilab.cruk.cam.ac.uk/mammaryGland/

Spatiotemporal expression is dynamically regulated throughout the adult mammary gland cycle (fig2a,b). In nulliparous mice, *Wnt7b* is expressed in luminal progenitor cells, as well as in more differentiated, hormone-sensing luminal progeny (fig2b,c). During gestation and lactation *Wnt7b* expression is switched off, but it reemerges post-involution (fig2b,c). Thus, it is exclusively expressed in the ‘resting’ state, be it nulliparous or post-involution. Of note, although the luminal progenitor population itself re-appears post-involution, *Wnt7b* expression is lost in this population, becoming restricted to the hormone-sensing luminal lineage post-pregnancy (fig2b,c).

From these analyses we would conclude that *Wnt7b* is expressed exclusively in the luminal compartment in the nulliparous mammary gland, is lost during pregnancy, and is re-established post-involution (fig3). Indeed, this is supported by other studies showing that *Wnt7b* is expressed in the virgin mammary gland, but drops at E12.5 of pregnancy to undetectable levels [7]. This underscores the validity of this approach and illustrates the usefulness of interactive *in silico* tools to determine spatiotemporal patterns of *in vivo* gene expression.

**Figure 3.**
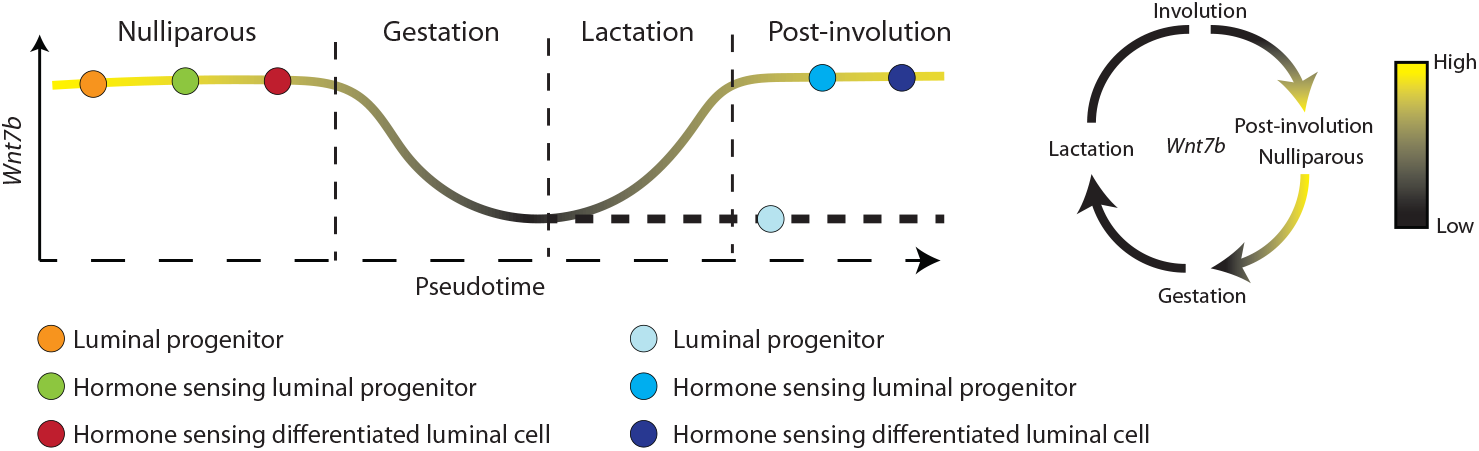
Graphic summary of mouse *Wnt7b* expression dynamics based on the scRNAseq data from Figure 1 and Figure 2. Drawn by the authors.

## Identifying putative regulatory elements

Little is known about the molecular signals and cis-regulatory elements that control mouse *Wnt7b* or human *WNT7B* gene expression. In ER-/HER2+ breast tumors, *WNT7B* was shown to be a direct transcriptional target of the androgen receptor (AR) [17] and predicted to be regulated by Nuclear respiratory factor 1 (NRF1) [18]. Although *Wnt7b* is also expressed in hormone-responsive cells (fig2 and [13]), at present there is no experimental evidence to support that it is regulated by steroid hormones, in particular progesterone [19]. *Wnt7b* expression is not limited to the mammary gland, however. It is required for lung [20,21], and kidney development [22] to name but a few and can therefore be regulated by a myriad of signals.

One way to gain understanding into tissue-specific gene expression, is to identify cis-acting enhancer elements. Using ChIPseq analysis, a recent study predicted 440 mammary-specific super-enhancers [23]. Super-enhancers can be classified as dense clusters of transcriptional enhancers that are likely to control genes important for cell type specification [23-25]. Only one of these was followed up in more detail in that particular study. However, a supplementary file listing all 440 of these putative regulatory elements is available. We were particularly intrigued by a sequence that spans more than 24 kb on chromosome 15 (published mm9 coordinates chr15: 85475778-85500063, mm10 coordinates chr15: 85645348-85669633), which was assigned as a putative regulator of the nearest gene: *Wnt7b* (fig4a). While it is common to do so, linear proximity alone is not an accurate measure for functional interaction between an enhancer and its putative target gene [24,25]. Other genes in this region – including two miRNAs (*Mirlet7c-2/Mirlet7b*) and a protein coding gene (*Ppara*) – might also be regulated by this particular super-enhancer. A region on the edge of this super-enhancer (mm9 coordinates chr15:85473689-85478592, published mm10 coordinates chr15: 85643259–85648162) was recently indeed associated with *Wnt7b*, albeit not in the mammary gland but in a mouse model for hair-follicle derived skin tumors, and based on strain-specific polymorphisms rather than on having been shown to directly regulate *Wnt7b* expression [14]. These results show that association of this super-enhancer with *Wnt7b* in the mammary gland is worthy of follow-up analysis.

**Figure 4.**
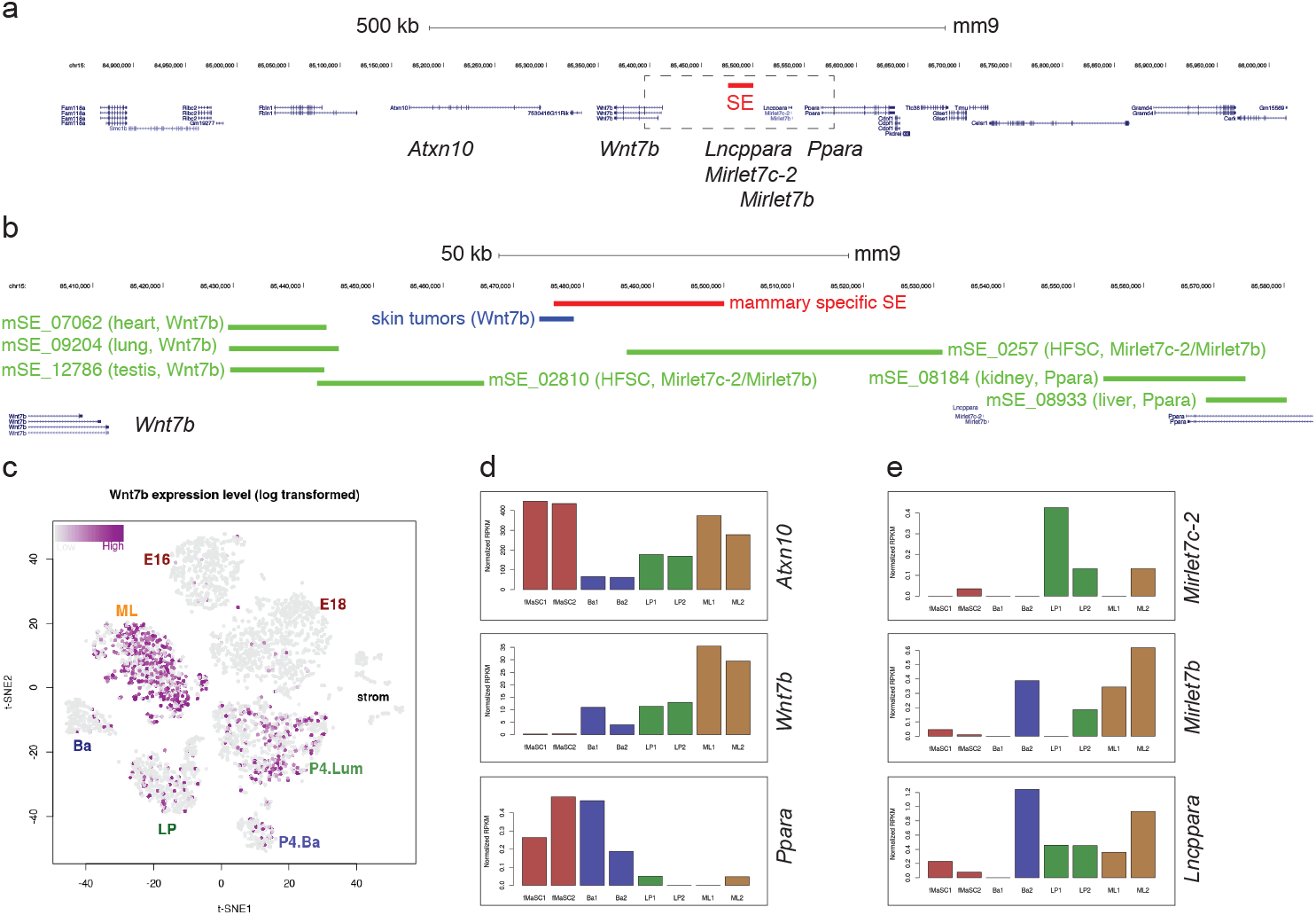
Overview of super-enhancers assigned to genes in the vicinity of *Wnt7b*. A) Location of a mammary-specific superenhancer (SE, in red) on mouse chromosome 15. Scale bar is 500 kb. B) Close up of the region boxed in A. The mammary specific SE is shown in red. A *Wnt7b* associated regulatory region in skin tumors is highlighted in blue. Other super-enhancers in this genomic region, listed in Superdb, are depicted in green. The tissue of origin in which they were identified and the genes to which they have been associated based on proximity rules are indicated. HFSC= hair follicle stem cells. Scale bar is 50 kb. C) tSNE plot of single cell *Wnt7b* expression from FACS-based scRNAseq data from [31]. D) Gene expression of annotated genes in the vicinity of the mammary gland specific SE for all epithelial mammary gland subpopulations. E) Expression of putative non-coding RNAs in the vicinity of the mammary gland specific SE for all epithelial mammary gland subpopulations. D-E show normalized RPKM values. Plots for C-E were generated at https://wahl-lab-salk.shinyapps.io/Mammary_snATAC/

The term “super-enhancer” is used to define a larger chromatin area that contains clusters of smaller, individual enhancers and that is enriched for active chromatin marks (e.g. H3K27ac) or occupied by transcriptional activators (e.g. MED1) and master regulatory transcription factors (e.g. STAT5A) [23,26,27]. More than 80,000 super-enhancers (combined numbers for the mouse and human genome) can be accessed through the online dbSuper database [28]. An updated version of the Super Enhancer Archive (SEA 3.0) provides another entry point [29], but this database was unfortunately offline when we were drafting this manuscript (Supplementary Table 1).

A first screen of the dbSuper database shows the tissue-specificity of superenhancers: a putative *Wnt7b* super-enhancer has also been identified in the murine heart, lung and testis. However, this sequence does not overlap with the mammary-specific super-enhancer described by Shin et al. [23]. Instead, the dbSuper database predicts this particular location to contain two super-enhancers, identified in hair follicle stem cells, linked to *Mirlet7c-2/Mirlet7b* [30]. Additional super-enhancers in this region, identified in the kidney and the liver, are tentatively associated with *Ppara* (fig4b). It should be noted that also in dbSuper, super-enhancers and their associated genes are linked based on a simply proximity rule to the nearest transcriptional start site (TSS) [28]. Out of the genes located in this ∼500 kb area on chromosome 15, only *Atxn10* and *Wnt7b* show prominent expression in one or more mammary gland cell subpopulations, although *Ppara, Mirlet7c-2/b* and a non-coding RNA, *Lncppara*, may be differentially expressed at low levels (fig4c-e).

## Determining the boundaries of the *Wnt7b* regulatory domain

In recent years, it has become generally accepted that regulatory elements control target gene expression within the confines of larger, structurally ordered regions of the chromatin known as topologically associating domains (TADs) [32]. Specific DNA sequences (i.e. regulatory elements and their target genes) are much more likely to interact within a TAD, than across a TAD boundary. A logical next step in exploring the potential regulation of *Wnt7b* by the aforementioned mammary-specific super-enhancer would therefore be to determine the boundaries of the *Wnt7b* TAD.

We used the 3D Genome Browser (Box 2) to visualize TAD predictions of the *Wnt7b* locus using publicly available Hi-C datasets [33]. In this browser, TAD boundary predictions are calculated according to the so-called directionality index, which is a method that looks at the degree of up- and downstream interaction bias for DNA regions [34]. It was noted that DNA regions at the periphery of TADs are highly biased in their direction of interaction. Upstream regions in a TAD are highly biased towards interacting with downstream regions and vice versa. Using this directional bias, the boundaries of adjacent TADs can be predicted. Their coordinates are provided by the 3D Genome Browser, which also includes an intuitive visual reference (fig5).

**Figure 5.**
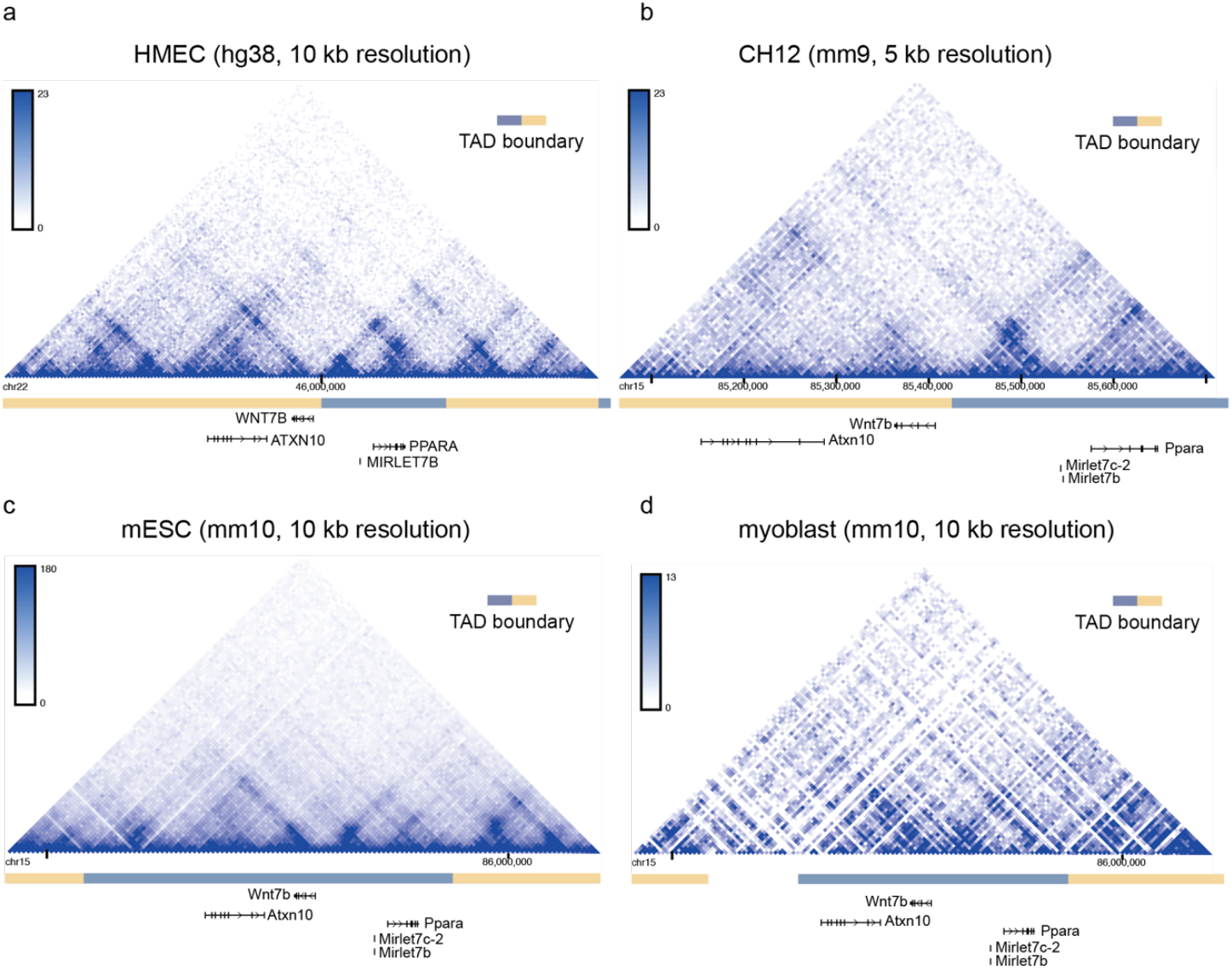
TAD boundary prediction using the 3D Genome Browser. In the depicted triangle, the physical interaction frequency of DNA regions is represented by the color intensity. A dark blue spot can be observed that connects the TAD boundaries of the predicted *Wnt7b* TAD, indicating that these genomic regions were found to frequently interact in this Hi-C dataset. Alternating beige and grey blocks (depicted in between the chromosome coordinates and the genes) depict individual TAD predictions. A) Hi-C data from human mammary epithelial cells, HMEC [35]. B) Hi-C data from mouse lymphoma cells, CH12 [35]. C) Hi-C data from mouse embryonic stem cells, mESC [38]. D) Hi-C data from mouse myoblasts [39]. Plots were generated at http://promoter.bx.psu.edu/hi-c/

**Box 2: Chromatin conformation capture Hi-C data**

*http://promoter.bx.psu.edu/hi-c/*

*(built by the Yue lab, described in Wang et al. (2018), The 3D Genome Browser: a web-based browser for visualizing 3D genome organization and long-range chromatin interactions* [33])

The 3D Genome Browser compiles published Hi-C and capture Hi-C datasets from both mouse and human cell lines or tissues, including HMEC. Chromatin conformation data from the locus of a gene or location of interest can either be displayed as a Hi-C heatmap or as a virtual 4C (with the location of interest as viewpoint). Where applicable, it will predict the boundaries of local TADs based on the provided dataset. In the upper menu bar are several options for visualizing data: In “HiC” you can visualize the data from different papers/datasets. In “Compare HiC” you can compare TADs from two different datasets. The coordinates of different TADs can also be downloaded in text file format for hg19, hg38, mm9 and mm10.

Only one mammary-specific Hi-C dataset is currently available, derived from human mammary epithelial cells (HMEC) [35]. However, TADs have been reported to be stable across cell types and even species [34,36]. Although not all TAD boundaries are equally stable [37], TAD organization can therefore also be investigated using Hi-C datasets generated from a different tissue as input.

According to this analysis, the *Wnt7b* TAD boundary lies immediately upstream of the *Wnt7b* TSS in both HMECs and mouse lymphoma cells (fig5a,b). This would imply that the mammary-specific super-enhancer identified by Shin et al. lies outside of the predicted *Wnt7b* TAD, which makes it less likely that this particular super-enhancer directly regulates the expression of *Wnt7b*. However, in other Hi-C datasets this TAD boundary is less well defined (fig5c,d).

## Discovering novel regulatory interactions

To gain a better understanding of how the spatiotemporal expression of *Wnt7b* is regulated in the adult mammary gland, we can start by probing the epigenetic state of the *Wnt7b* locus in an R shiny app published by the Wahl lab (Box 3). This tool not only allows chromatin accessibility and relevant histone modifications to be examined, but also can be used to make predictions about specific promoters and their regulatory sequences of interest. An attractive graphical interface allows intuitive interpretation of the data (fig6).

**Figure 6.**
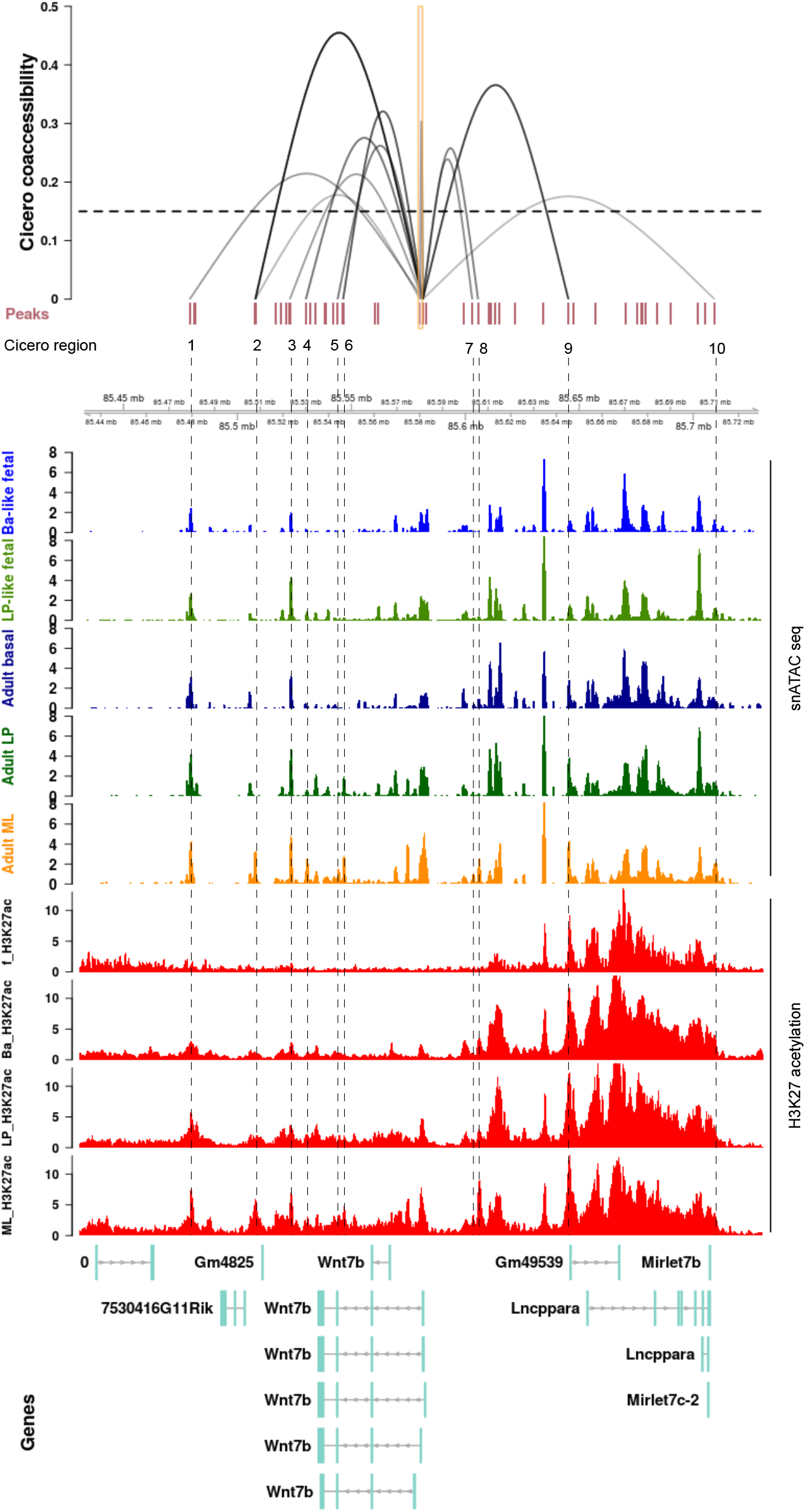
Overview of chromatin accessibility, genomic interactions and the epigenetic status of the *Wnt7b* locus. Top) Regions identified by Cicero as having a higher co-accessibility score than 0.15 are displayed as interacting loops with the *Wnt7b* promoter. The height of the loop indicated the corresponding Cicero score. The viewpoint size at the promoter is a stretch of 1000 bp. Middle) The 4 snATAC tracks display the aggregated snATAC signal from [40] at the *Wnt7b* locus for each epithelial mammary gland subpopulation. Ba-like: Basal-like, LP: Luminal Progenitor, ML: Mature Luminal. Bottom) The 4 H3K27 acetylation tracks display bulk ChIPseq of FACS sorted cells from [41] at the *Wnt7b* locus for each epithelial mammary gland subpopulation. ML_H3K27ac: Mature Luminal, LP_H3K27ac: Luminal Progenitor, Ba_H3K27ac: Basal, f_H3K27ac: fetal. All the data is aligned to mm10 and displayed in a window size of 300 kb. Note that *Wnt7b* is oriented in the reverse orientation (i.e. expressed from the minus strand). Regulatory regions 1-6 to the left of *Wnt7b* are therefore downstream of the promoter and regions 7-10 to the right of *Wnt7b* are upstream. Distal elements that are predicted to interact with the *Wnt7b* promoter and alter their epigenetic status in accordance to *Wnt7b* expression can have the potential to be involved in the spatial temporal regulation of Wnt7b in the mammary gland and therefore warrant further investigation. The shiny app offers an intuitive and interactive visual tool to quickly compare numerous epigenetic features, and identify novel regions of interest. It should be noted that no statistical analysis or specific coordinates are provided, although these are available in supplementary data and the GEO accession file. Hence, it serves as an excellent hypothesis generating tool that requires further validation either by *in silico* analysis or experimentation.

**Box 3: Probing chromatin accessibility and epigenetic interactions**

*https://wahl-lab-salk.shinyapps.io/Mammary_snATAC/*

*(Chung et al*., *2019, Cell Reports* [40] *& Dravis et al*., *2018, Cancer Cell* [41]).

The R shiny app published by the Wahl lab combines bulk RNAseq and H3K27 acetylation ChIPseq data from [41] with single-nucleus ATACseq (snATACseq) and scRNAseq data from [40] in an online web interface that allows its users to investigate numerous (epi)-genetic feature in fetal mammary stem cells (E18 fMaSCs), basal, luminal progenitor and mature luminal cells. This allows researchers to investigate single cell expression & chromatin state (accessibility in the case of snATACseq and active enhancer marks in the case of H3K27Ac ChIPseq) of their gene of interest, and to follow expression of the gene along a pseudotime trajectory. Moreover, if this gene is a transcription factor, its activity can be predicted for each subpopulation based on motif enrichment in open chromatin regions from snATACseq data. Lastly, based on co-accessibility of distal sites and promoter regions in single cells promoter-enhancer interactions for the gene of interest can be predicted using the so-called Cicero algorithm [42], and concurrently displayed with chromatin accessibility scores and H3K27Ac from aggregate snATACseq and bulk ChIPseq data. In the online tool, Cicero makes predictions in a region of max. 300 kb (with 150 kb upstream and 150 kb downstream of the viewpoints).

If we focus our attention on the *Wnt7b* promoter and gene region (i.e. the center portion of fig6), snATACseq reveals that the chromatin is relatively accessible in all mammary cell type subpopulations irrespective of *Wnt7b* gene expression levels (fig6, top 5 rows). In contrast, H3K27ac of the *Wnt7b* promoter and gene region is exclusively enriched in the luminal compartment (fig6, bottom 4 rows in red). This suggests that *Wnt7b* is ‘primed’ and open in all epithelial cells in the mammary gland, but its potential for increased gene expression is only realized in the luminal compartment where the chromatin displays the proper histone acetylation marks.

Combining the Cicero algorithm (see Box 3) with snATACseq data, this online tool can also be used to infer co-accessibility of distal sites and the promoter of their putative genes in individual cells. In this manner, Cicero can predict cis-regulatory elements that would be able to interact with the *Wnt7b* promoter *in vivo*. At a co-accessibility threshold of 0.15, Cicero identifies 10 regions within 150 kb up-or downstream of the viewpoint that interact with the promoter of *Wnt7b*. Of these, 4 are located upstream of *Wnt7b* in an area dense with H3K27ac that encompasses, but extends beyond, the super-enhancer region, and 6 are located downstream of *Wnt7b* (fig6).

The interacting regions depicted to the left of the *Wnt7b* promoter (regions 1-6, located 3’ distal to the TSS) all fall within in the predicted *Wnt7b* TAD (compare fig5,6). These distal sites are either somewhat enriched for chromatin accessibility or H3K27ac, or a combination of both epigenetic features, in adult luminal progenitor and mature luminal cells compared to the adult basal subpopulation (fig6,7). The 4 regions downstream of *Wnt7b* (7-10) do not display evident changes in chromatin accessibility or H3K27ac when luminal cells are compared to the basal compartment, except for region 8 (fig6,7). Note that the distance between region 9 and 10 spans more than 60 kb, which is considerably larger than the reported size of the mammary-specific super-enhancer. This entire stretch of 60 kb shows characteristic marks of active and open chromatin, suggesting that a much larger collection of regulatory elements may exist in this area (fig6).

**Figure 7.**
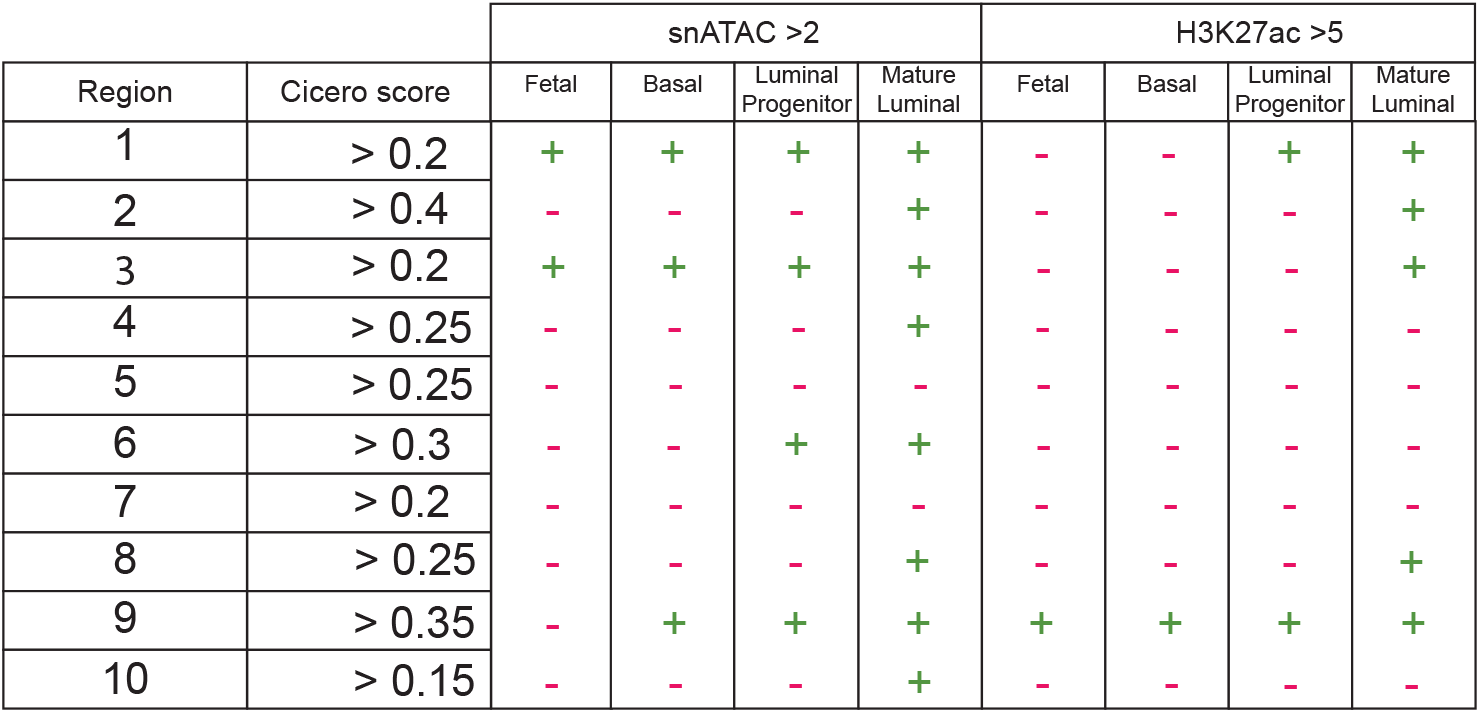
Summary of the epigenetic features of each region identified by Cicero as depicted in Figure 6. Somewhat high(er) levels of snATAC seq signal are defined above a cut-off of 2 and H3K27 acetylation above a cutoff of 5. Note that the tool does not offer any statistical analysis, and therefore cut-offs were user-defined compared to the total signal in the 300 kb window. They should thus be considered reasonable, but relatively arbitrary and worthy of more in-depth investigation.

**Box 4: Looking for evolutionary conservation**

http://ecrbrowser.dcode.org

*(Ovcharenko et al*., *2004, Nucleic Acids Research* [43]).

This web-based tool enables access to pairwise alignments for the genomes of 13 species and visualizes evolutionary conserved regions (ECRs) in a graphical interface. Users can set their own parameters to select regions with a desired cut-off (in figure 8: >85% sequence identity over >200 bp). Sequences that are conserved within these chosen parameters are represented as colored peaks. Conservation between species is shown relative to a base genome of choice (in figure 8: Hg19). Sequence information from the UCSC Genome Browser (http://genome.ucsc.edu/) [44] can be extracted when selecting the DNA region of interest.

Genome assemblies used in the ECR browser: human: Hg19, Tetraodon: tetNig1, frog: xenTro3, fugu: fr3, zebrafish: danRer7, chicken: galGal3, opossum: monDom5, rat: rn4, mouse: mm10, cow: bosTau6, dog: canFam2, chimpanzee: panTro3, rhesus macaque: rheMac2.

**Figure 8.**
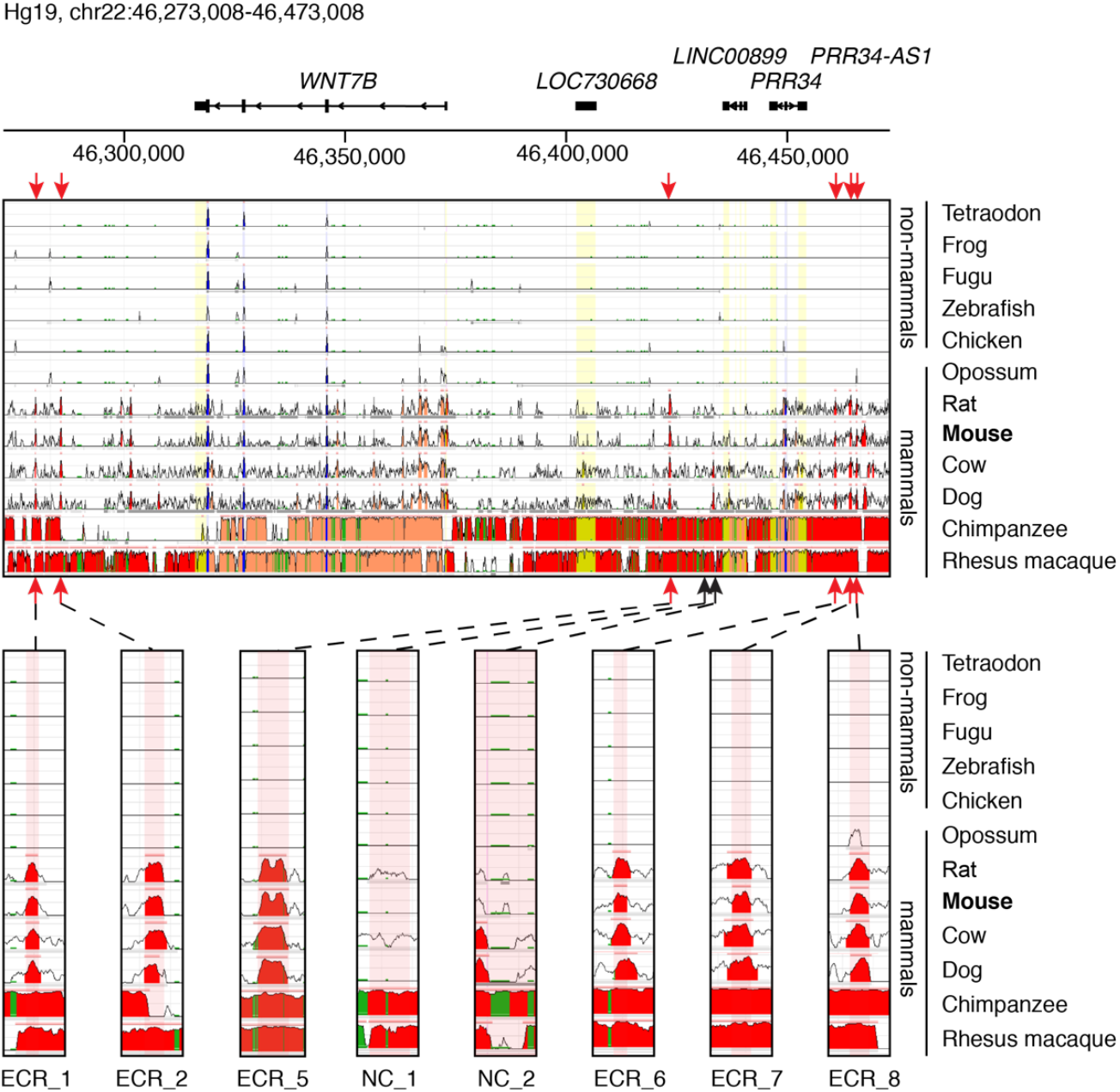
Selection of candidate enhancers based on sequence conservation. A 200 kb region of the human Hg19 genome assembly, 100 kb up-and downstream of *WNT7B* TSS. In each track the sequence conservation between Hg19 and one of 12 vertebrates is shown. Different colors indicate the following: Red = intergenic, salmon = intragenic, yellow = UTRs, blue = coding sequences, green = transposons and simple repeats. Locations of putative candidate *Wnt7b* enhancers are indicated by red arrows. Lower panels show zoomed in regions of 1000 bp where the candidate enhancers are located. The red shade over the tracks represent the chosen candidate enhancer region. Examples of sequences that are conserved in mammals, but not in non-mammalian vertebrates are shown (ECR_1, ECR_2, ECR_5, ECR_6, ECR_7 and ECR_8) alongside two examples of a non-conserved region (NC_1, NC_2). Parameters used: >85% sequence identity over >200 bp.

## Exploring conservation of putative regulatory enhancer sequences

In previous studies, highly conserved sequences were associated with developmental and transcriptional regulators [45–51]. Given the fundamental role of Wnt signaling not only in vertebrate development [52], but also specifically in mammary gland development and maintenance [53–55], focusing on conserved sequences could be another criteria for the selection of candidate *Wnt7b* enhancers. To identify conserved regions in the vicinity of *Wnt7b*, we used the evolutionary conserved region (ECR) browser (Box 4).

Often, conservation is scored across vertebrate species. However, in an attempt to identify regions that are specifically conserved in mammals, we specifically selected candidate sequences in a region of ∼100 kb up-and downstream of the *Wnt7b* TSS that are conserved across mammalian, but not necessarily in non-mammalian vertebrate species available in the ECR browser (fig8).

## A working model for follow-up studies

Of course, none of these approaches (sequence conservation, histone modification, transcription factor ChIPseq), either by themselves or in combination, are sufficient to definitively link any of these putative regulatory elements to *Wnt7b*. This requires further experimental validation and specific follow up. However, as a prediction tool these combined analyses provide an excellent starting point for dissecting this super-enhancer in more detail. If we put all of the different pieces of information together (fig9), we can draft some hypotheses regarding the regulation of *Wnt7b* expression in the mammary gland.

**Figure 9.**
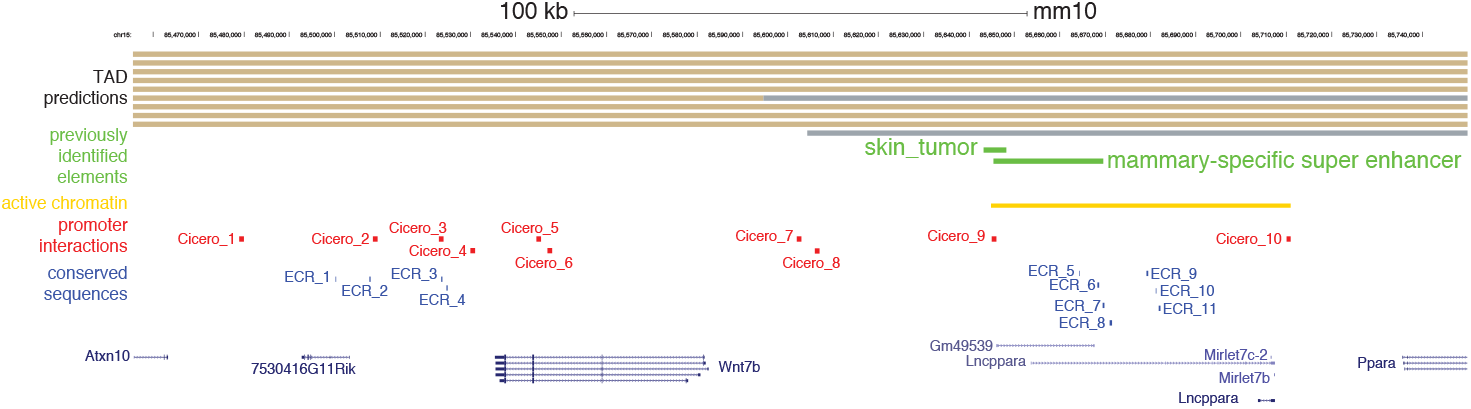
Integration of the insights obtained from the different analyses. Beige and grey bars at the top represent neighboring TADs as predicted in cortex (liftover from mm9), mESC (liftover from mm9), neurons (mm10), mESC (mm10), NPC (mm10), CH12 (liftover from mm9), cortical neurons (mm10), myoblast (mm10), G1E-ER4 (mm10) and HMEC (liftover from human). Previously identified (super) enhancer elements are depicted in green, the region of active chromatin identified in the Cicero analysis is depicted in yellow, promoter interactions predicted by Cicero are depicted in red and conserved sequences identified in the ECR browser are depicted in blue. Coding and non-coding genes are shown at the bottom for reference.

First, we propose that in mammary epithelial cells the proposed TAD boundary immediately upstream of *Wnt7b* (fig5) is not very stable, given that the Cicero algorithm predicts four interactions between the *Wnt7b* promoter and regions to the right of this presumed TAD boundary (i.e. regions 7-10 in fig6). Of note, two of these interactions (Cicero regions 7 and 8) occur in the direct vicinity of this presumed TAD boundary. The other two interactions (Cicero regions 9 and 10) border a large area of active chromatin, which extends beyond the super-enhancer region previously identified by Shin et al. [23]. It has not escaped our attention that this 60 kb area harbors an annotated lncRNA (*Lncppara*) and two microRNAs, *MirLet7b/MirLet7c-2*, which are broadly expressed and implicated in cancer formation [56–58]. Moreover, this region also contains multiple conserved sequences that could represent functional enhancer elements (including ECR_6, ECR_7 and ECR_8 from fig8).

Second, if we do take the TAD boundary prediction into account, it may be wise to prioritize the interactions that occur between *Wnt7b* and more downstream sequences (i.e. regions 1-6 in fig6). Although the coordinates from the Cicero prediction algorithm deserve further scrutiny of the original datasets, these downstream interacting regions also lie in close vicinity to conserved sequence elements.

Third, in combination with the expression data analysis (fig1,2), the published literature and the active enhancer marks (fig6,7), we can make a further prioritization of putative *Wnt7b* enhancer sequences that are worthy of experimental validation and follow up. In this case, region 2 is particularly interesting as is has the highest Cicero score and displays both differential chromatin accessibility and H3K27 acetylation in the luminal compartment.

To summarize, by using publicly available online tools we assessed the genomic conformation of the *Wnt7b* locus, and how this relates to the previously identified putative *Wnt7b* super enhancer. By examining the epigenetic status of the *Wnt7b* locus more closely, we noticed that although the *Wnt7b* promoter is predicted to interact with the super-enhancer region, this is likely not cell type specific as both chromatin accessibility and H3K27ac do not change between the basal and luminal lineages in this region. However, regions downstream of *Wnt7b* do change their epigenetic status in accordance to *Wnt7b* gene expression and are also predicted to interact with the *Wnt7b* promoter. This entire area would be worthy of experimental follow up to definitively associate specific regulatory elements with *Wnt7b* and/or other nearby genes – in particular the miRNAs and *Ppara*.

## Discussion

Using publicly available genome wide datasets and accessible online tools, we have identified several regions that might play a role in the regulation of spatiotemporal expression of *Wnt7b* in the mouse mammary gland is regulated. Our main goal was to show the reader how these findings provide additional information for future investigations. However, we also want to use this opportunity to highlight and stress the added value of making large datasets available to a wide audience through interactive online tools. We thank our colleagues who invest their resources to do so. At the same time, we call for joint efforts from our community to ensure that the repertoire of tools as well as of accessible datasets continues to grow and remains of high quality and value to investigators worldwide. As others have undoubtedly noticed, mammary gland and breast tissue datasets are often notoriously absent from public, large-scale -omics efforts. Generating and curating additional genome wide datasets (e.g. Hi-C and others) for both epithelial and stromal cells of multiple species, including mouse and human, would be a tremendous resource for our community as a whole. The careful generation of such datasets in combination with user-friendly online tools provide a valuable resource for researchers, and could in the long run also help to reduce animal experimentation. Certain features will enhance the user experience and promote the wide use of such tools, including the ability to export high resolution graphs (ideally allowing further customization, e.g. PDF format as offered by [15,33]) and the ability to easily download specific sequences or genome coordinates (as offered by [33,43]). Given the challenges associated with keeping these databases up to date and operating smoothly, international and consortium efforts that provide sufficient support infrastructure may, in the long term, prove to be essential in this regard.

Here we have shown how the combined use of different online tools can be applied to generate novel hypotheses. Of course, the same tools can also be used to complement existing projects by providing additional data. Ideally, in the not too near future, researchers will have a broad compendium of resources available to them that are of such high quality that they will allow *in vivo* analyses to be performed *in silico*, thereby bringing such genome-wide analyses within reach of all scientists. This will only be possible, however, if sufficient tissue-specific datasets can be accessed. Especially in the case of the mammary gland, great care should be taken to include different timepoints to cover both embryonic and postnatal developmental stages, as well as the entire gestational cycle. Here, biological and computational expertise will continually need to go hand in hand to ensure that such online tools can meet the demands of the scientific questions that are being asked.

## Author contributions

Conceptualization: YBCvdG, RvA; Methodology/Experiment design: YBCvdG, NH, RvA; Investigation/Data acquisition: YBCvdG, NH, RvA; Formal analysis/Data interpretation: YBCvdG, NH, RvA; Writing – original draft: YBCvdG, NH, RvA; Writing – revision and editing: YBCvdG, NH, RvA; Visualization: YBCvdG, NH, RvA; Supervision: RvA; Approval final manuscript: YBCvdG, NH, RvA; Project administration/Stewardship: YBCvdG, RvA; Funding acquisition: RvA.

---

**Supplementary Table S1.**
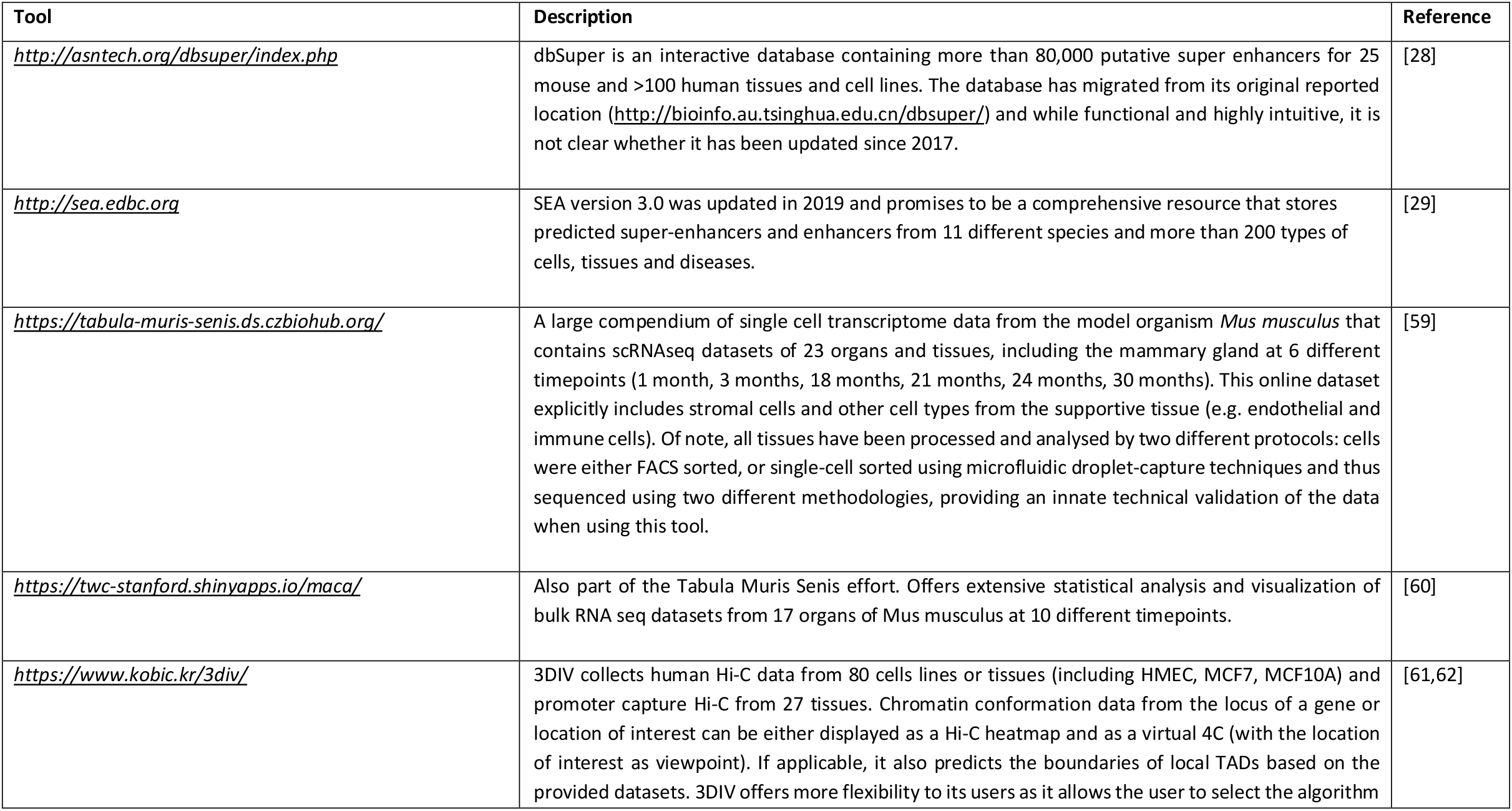
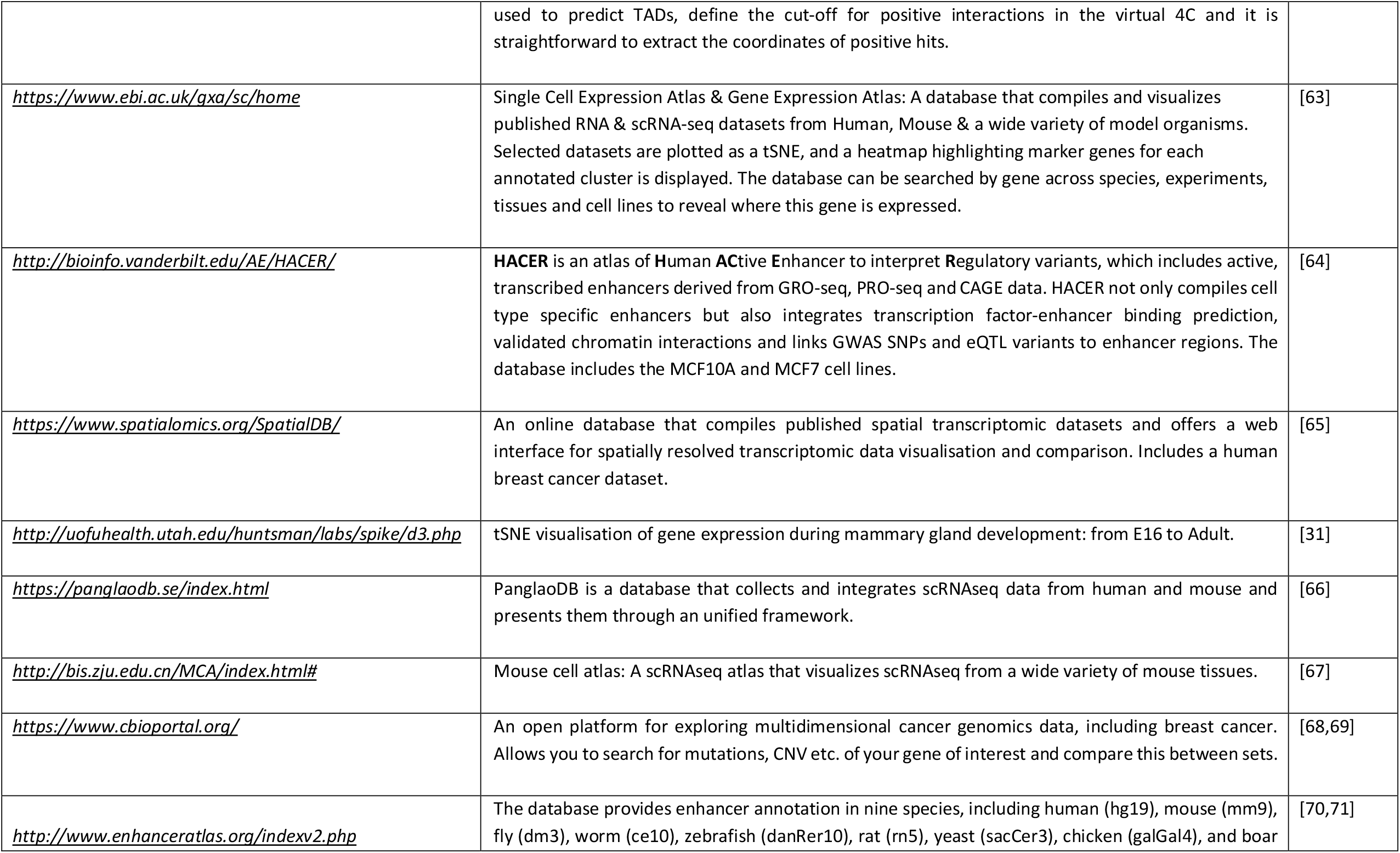
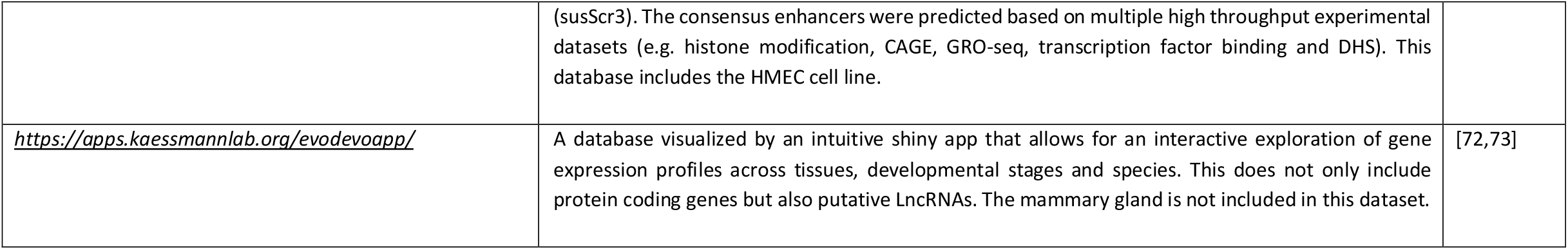
Compilation of publicly available online tools that are outside the scope of the current case study. These tools are not specific for mammary gland biology and/or do not always include mammary gland datasets.

